# To increase trust, change the social design behind aggregated biodiversity data

**DOI:** 10.1101/157214

**Authors:** Nico M. Franz, Beckett W. Sterner

## Abstract

Growing concerns about the quality of aggregated biodiversity data are lowering trust in large-scale data networks. Aggregators frequently respond to quality concerns by recommending that biologists work with original data providers to correct errors “at the source”. We show that this strategy falls systematically short of a full diagnosis of the underlying causes of distrust. In particular, trust in an aggregator is not just a feature of the data signal quality provided by the sources to the aggregator, but also a consequence of the social design of the aggregation process and the resulting power balance between individual data contributors and aggregators. The latter have created an accountability gap by downplaying the authorship and significance of the taxonomic hierarchies - frequently called “backbones” - they generate, and which are in effect novel classification theories that operate at the core of data-structuring process. The Darwin Core standard for sharing occurrence records plays an under-appreciated role in maintaining the accountability gap, because this standard lacks the syntactic structure needed to preserve the taxonomic coherence of data packages submitted for aggregation, potentially leading to inferences that no individual source would support. Since high-quality data packages can mirror competing and conflicting classifications, i.e., unsettled systematic research, this plurality must be accommodated in the design of biodiversity data integration. Looking forward, a key directive is to develop new technical pathways and social incentives for experts to contribute directly to the validation of taxonomically coherent data packages as part of a greater, trustworthy aggregation process.

## INTRODUCTION

Many fundamental problems in biology rely on data about the locations and traits of naturally occurring organisms classified according to taxonomic categories. These occurrence records are often tied to vouchered observations or specimen depositions with provenance from natural history collections, legacy publications, data repositories, and ongoing inventories (1). Numerous projects are bringing occurrence records into the realm of modern data science (2-6). This process involves multiple levels of data creation and aggregation to enable the synthesis of biodiversity data trends at greater scales (7-9). However, there is a widespread sense that aggregated biodiversity data should be used with caution because they are frequently of insufficient quality to support reliable inferences.

Over two dozen studies have found quality shortcomings in aggregated occurrence datasets (9-37). The growing concerns are leading to reduced trust in the aggregators’ data, and thereby also reduce the data's scientific use and societal impact (38).

Biodiversity data aggregators frequently respond to quality concerns by recommending that biologists work with the original providers of the data to correct errors *at the source* (19). This argument is exemplified by Sikes et al.’s response (33: 149) to Ferro and Flick (26): “[W]e have heard some taxonomists state they do not want to share data with GIBF [the Global Biodiversity Information Facility] because they distrust the quality of the data in GBIF. This latter point seems illogical. The data in GBIF are the data from the museums that provide data. If the data in GBIF are not to be trusted then neither are the data in the source museums. It thus seems illogical to be pro-museum and anti-GBIF”.

We show that this strategy of allocating responsibilities for data quality falls systematically short of a full diagnosis of the underlying causes, because it fails to make sense of an important way in which many biologists have come to mistrust aggregators. The distrust held for biodiversity data aggregators *is* justified once we recognize that the aggregation process is designed in a way that holds no one accountable for the accuracy of the taxonomic hierarchies operating at the core of their data-structuring process.

On the surface, aggregators appear to recognize the need for such accountability (19: 73): “What ideally is needed is an environment created by agencies such as GBIF and ALA [the Atlas of Living Australia] that *efficiently* enables” the exposure, discussion, and correction of errors “directly in *all relevant locations*”. The authors acknowledge that “[n]o such environment currently exists”, and “[p]rogress will be limited while the underlying culture of data publishing and data management does not support stable, long-term reference to each data record and community-based curation of those data in a way that ensures that each act of correcting *any aspect of any data element* is not lost but contributes to the development of a global digital biodiversity knowledgebase” (19: 73-74; emphasis added).

We argue that in practice aggregators have made fundamental design choices that lower trust in the aggregation process by excluding taxonomic conflict as an aspect of biodiversity knowledge worthy of being preserved. Looking ahead, we need new solutions capable of handling common situations in which authors endorse and reconcile incongruent systematic hypotheses. Based on this diagnosis, we recommend shifting the balance of power to license trustworthy data packages away from biodiversity aggregators and towards individual expert authors.

## No simple diagnosis for data quality deficiencies

Database systems constrain as well as enable science. They have the power to channel research; e.g. by making some questions harder to ask, by entrenching certain presuppositions into their design, and by placing new burdens on participation on the research community (39-44). Implementing data aggregation systems therefore places one in a position of responsibility for the larger community – a status that is at times novel for aggregators who by training are more focused on the domain-specific aspects of science than social engineering. Abstract notions of responsibility become concrete, however, when they intersect with the ways that an aggregation system handles the process of correcting data quality deficiencies.

We take the presence of a growing corpus of critical studies, such as those cited above in the Introduction, as evidence that researchers are becoming increasingly cautious about relying on aggregators. Other biologists are bound to notice and may be deterred by the costs required to carry out the necessary additional quality vetting. That said, the helpful term “fitness-for-use” (16) reminds us that certain deficiencies can be acceptable to researchers whose inference needs are robust to them. Many biodiversity data packages and analytical needs will match up well in this sense. Nonetheless, providing data that are just clean enough for coarse-grained biodiversity inferences is not a way to deal with the root issues affecting trust.

We further believe that all parties – from individual record authors to high-level aggregators – have an interest in precise diagnoses of where the responsibilities for specific trust deficiencies lie. Aggregators are not helped in the long run by a diagnosis that does not lead each party to play an appropriate role in overcoming these issues.

To review: a common aggregation path for an occurrence record includes (i) individuals recording the original data and metadata at the time of collection; (ii) transcribing the record from field notes into the source collection’s local database; and (iii) applying further data and transformations – such as georeferencing and taxonomic identification – to comply with locally accepted conventions. Often, (iv) the collection will transfer the record to a regional, mid-level aggregator, which may then (v) transmit the information again to a higher-level aggregrator. Alterations of ‘the record’ can occur at any stage along this provenance chain, and have the potential to affect the identity of the record and its empirical signal.

Responsibilities for generating and repairing quality issues are not uniformly distributed across the data aggregation chain. Accordingly, the question of who is responsible for creating and fixing deficiencies cannot have a simple, univocal answer. Correcting a false collecting date would typically be the responsibility of individuals with knowledge of the original collecting event. Dealing with repeated duplication of records *through* aggregation, in turn, is a task better suited for aggregators. Two examples illustrate this point. First, in their response to Mesibov’s (22) audit of aggregated data on Australian millipedes, Belbin et al. (19) – representing the aggregator's perspective – provide a table with 44 automated quality checks performed by ALA (45). The five most frequent categories of error, respectively affecting 18.6–90.2% of the records, are all related to georeferencing. Second, Hjarding et al. (28), in their assessment of records of East African chameleons served by GBIF (46), conclude that 99.9% “used outdated taxonomy” that would have led to inadequate threat category assignments for eight taxa.

To extend the point about diagnostic precision further, Darwin Core (DwC), the prevailing global standard for sharing occurrence records, has seven main categories (1). Of these, Event, Location, and Taxon are the primary data blocks where insufficient quality will affect inferences of biodiversity distributions. How abundant and significant the shortcomings are will vary from case to case.

## Importance of trust for sustained use

Trust is a complex and context-sensitive concept (47-51). Our use of the idea will be anchored in two core assumptions. First, trust is a dependence relation between a person or organization and another person or organization: the first agent depends on the second one to do something important for it. An individual molecular phylogeneticist, for example, may rely on GenBank (52) to maintain an up-to-date collection of DNA sequences, because developing such a resource on her own would be cost-prohibitive and redundant. Second, a relation of dependence is elevated to being one of trust when the first agent cannot control or validate the second agent's actions. This might be because the first agent lacks the knowledge or skills to perform the relevant task, or because it would be too costly to check. Historically, when phylogeneticists published individual gene sequences or alignments in their research articles, one could actually check by unaided visual inspection whether a GenBank entry was correct (40, 53). Nowadays, few biologists are able to validate ‘directly’ (i.e., without informatics tools) whether a genome sequence was correctly assembled from next-generation sequencing data.

Trusting someone means being in a position of dependence, but not necessarily a defenseless one. Individuals often rely on higher-order signals about an agent’s reliability and competence, whose value does not require knowing what that agent knows or being there to see what the agent actually does (48,54-55). Talking to biologists who have worked with an aggregator's data in the past, for instance, is a common way to learn whether those data can be trusted as accurate and for what purposes. Thus, the critical studies cited above (9-37) are valuable in part because they move the experience of working with aggregated data into the public domain. Reading these articles is analogous to reading on-line reviews about a contractor's performance on past jobs.

It is at least as important to know what an agent does when a task goes wrong, particularly for complex tasks like building a house or a higher-level biodiversity data aggregation service. Trust, then, is tied to more than just the intrinsic accuracy of the data! It is never just about whether the data are right or wrong, in the eyes of one or another party. Instead, trust also depends critically on how the trusted agent responds to negative outcomes – whether due to epistemic disagreements, honest errors, unanticipated difficulties, or negligence (55-57).

More succinctly, trust is lowered when there is a mismatch between “responsibility” and “admission” – i.e., an accountability gap – in the context of data shortcomings. Transparency about ongoing progress or known problems is often crucial to cementing an ongoing relationship of trust, because it demonstrates that the trusted agent is willing to pay the cost of generating regular reports (58-59). For example, some aggregators are intentionally transparent about letting experts identify data deficiencies by allowing the flagging of records with perceived problems (45, 60).

## Matching accountability to responsibility

It is especially harmful if the trusted agent creates new problems in an area of central concern to those relying on it while refusing to be held accountable for correcting them. We will argue that this is the case with the generation of synthetic taxonomic classifications and phylogenies – so called “backbones” or “trees of life” – that prevail in many biodiversity data aggregation networks (2, 61-65). High-level choices in designing these environments routinely lead to the creation of novel hierarchical syntheses that re-structure the occurrence record data in scientifically significant ways (42). In doing so, aggregators compromise established conventions of sharing and recognizing taxonomic work.

It is pivotal, then, that aggregators have embraced a design paradigm that requires one hierarchy (at a time) to organize all occurrence data. The production of a unitary hierarchy is governed by feasibility constraints, such as computational costs and institutional limits on sharing access to information, rather than principles based in best systematic theory and practice. However, achieving a unitary hierarchy requires aggregators to eliminate taxonomic conflict between data input sources. This often results in a hierarchy that no longer corresponds to the view of any particular source: it becomes a synthesis nobody believes in. Biologists frequently regard the quality of these novel classification theories as deficient.

Responsibilities for issues with aggregator-created synthetic classifications are not rightfully owned by any data-providing source. In particular, a one-time fix (adjustment) of the synthesis to match one source classification fails to correct an underlying design flaw: to achieve trustworthy bodies of data for biodiversity research and conservation, we need to manage each body of data as a coherent whole, not just as an aggregate that is somehow supposed to maintain unity at any scale. Since high-quality data packages frequently mirror competing and conflicting classifications, i.e., unsettled systematic research, this plurality must be accommodated in the design of biodiversity data integration.

## Generation of novel systematic syntheses

We use the term “systematic syntheses” as an encompassing term for different ways of organizing biodiversity into hierarchies, ranging from Linnaean taxonomies to phylogenetic trees. Achieving synthesis is the explicit goal of aggregators such as GBIF (65), which assembles its classification from more than 50 sources which are themselves unitary systems for particular subdomains of life (61, 66-67).

The aim to achieve one natural hierarchy is perhaps as old as systematics, remains valid today, and need not be dissected here. Similarly, the broader feasibility and merit of generating backbones have been discussed before in the present context (2, 68-70). The main, likely uncontroversial conclusion, that we carry over from these exchanges is this: in all instances where alternative, lower-level input classifications (subtrees) are available, it is necessary to select one schema over the alternative(s) to create the synthesis. This process often involves input from socially sanctioned individuals and committees, or (increasingly) the use of computer algorithms that resolve conflicts according to programmed criteria (62, 71-74).

Regardless of whether achieved by committee or algorithm, systematic syntheses typically have unique histories of creation and – by virtue of preserving the choices made to resolve conflicts – include or exclude information about how life on earth is structured. In accordance with Leonelli (42), these syntheses constitute *novel* classification *theories.* “Some classificatory systems systematically and synthetically express, rather than simply affect, knowledge obtained through scientific research, and they do it in a way that (1) is unique, since such knowledge is not formalized anywhere else in the same way; (2) has huge influence on knowledge-making practices; and (3) enables experimenters to make sense of the results they obtain. […] Articulating knowledge that enables scientists to assess and value their results is an achievement that goes well beyond listing a set of commonly used assumptions as a basis for further inquiry. In the latter case, existing knowledge is applied to put a given set of items into some order; in the former, existing knowledge is transformed and developed so as to facilitate the conceptual analysis of data” (42: 344-345).

To give one example, the 2016 version of the GBIF taxonomy (65), which is largely but not fully congruent with Ruggiero et al. (75), recognizes 99 phyla in eight kingdoms. Of these, two phyla are in the Archaea sec. GBIF Secretariat (65) and 29 phyla are in the Bacteria sec. GBIF Secretariat (65). We use the “sec”. – according to – to specify the source’s name usage (30-31). Meanwhile, Hug et al.’s (76) “new view of the tree of life” recognizes 26 phyla of Archaea sec. Hug et al. (76) and 92 phyla of Bacteria sec. Hug et al. (76). While Hug et al.’s (76) hierarchy is more transparently advertised as an outcome of systematic inference than the GBIF taxonomy (65), both are novel and unique in terms of their systematic content. Using one over the other meets Leonelli’s (42) criteria, i.e., such choices will influence how knowledge is transformed and how data analyses are facilitated.

To give another example, primate taxonomy is currently experiencing a period of destabilization, where the definitional boundaries of primate species remain subject to disagreement in light of increasing amounts of data and alternative inference tools (31, 77-78). The GBIF taxonomy has an influential function in this context, because newly indexed records whose identified species-level names are not endorsed by the hierarchy are ‘matched’ to it in one of two ways (21): either (1) names recognized as invalid versions (synonyms) of valid names are replaced by those, or (2) names that are not recognized at all at the species level are represented only the next available higher rank (e.g. genus), with a “null value” at the species rank. In fairness, it is possible to retrieve the original identifications on-line for individual records (one by one), or to download these as a complete spreadsheet-based dataset. However, this is not the same as representing all source records in their original taxonomic configuration and through the aggregator’s user interface.

The act of modulating the taxonomic identity of an occurrence record is a form of scientific arbitration that differs from a mere “enabling” of that record for aggregation and retrieval. The act may challenge and overrule the original judgment of taxonomic validity at the same nomenclatural rank, or may altogether reject and change that rank assignment. Even if this is not the case, transformation of the original taxonomic identities of records to match the chosen hierarchy constrains these records to reflect the taxonomic judgments endorsed by the hierarchy.

## Role of the Darwin Core standard

The Darwin Core standard (1) plays an under-appreciated role in creating the accountability gap. Darwin Core competes in this context with another standard for exchanging biodiversity data: the Taxonomic Concept Transfer Schema (TCS; 79). The TCS has a more limited scope than DwC with regards to non-taxonomic meta-/data properties of occurrence records. Yet the TCS was specifically designed to represent and integrate change and conflict across systematic hierarchies (30-31, 79-81). One of the key TCS conventions is to manage “taxonomic concept labels”, i.e., to consistently refer to name usages *according to* (sec.) a particular source. This allows the assembly of multiple coherent taxonomic hierarchies, each of which may assign incongruent meanings to overlapping sets of names within one shared platform. The taxonomic concepts endorsed by each hierarchy can be articulated using the relationships of Region Connection Calculus (RCC-5), which are part of the TCS. Such an approach can yield logically consistent, multi-hierarchy reconciliation maps (alignments), without disrupting the perspective advocated by each data source (30-31, 82).

By processing biodiversity data primarily via DwC, aggregators buy into a model that fails to represent the sources’ data signals directly. Because DwC lacks the syntactic conventions needed to absorb and align conflicting taxonomic hierarchies, the choice to favor DwC-over TCS-based solutions – minimally as a complement that adds syntactic precision to DwC where needed – becomes a systemic constraint of the aggregation design.

Problems with relying just on DwC syntax are further amplified by widespread implementation practices. For instance, while DwC permits the use of taxonomic concept labels at the species level, the standard does not forbid leaving the “dwc: nameAccordingTo” category empty. Because enforcing such labels is optional in DwC, doing so now becomes a social responsibility. In practice, the most occurrence records that participate in global aggregation are syntactically under-specified at the species level.

Conversely, DwC does not require filling in higher-level taxonomic names (genus, etc.) for occurrence records. Yet in this case the option to do so *should* be ignored, because above the species-level DwC syntax does not permit using taxonomic concept labels at all. Hence the DwC-permissible higher-level name usages are necessarily under-specified. In either case, these higher-level names need not travel along with every occurrence record. The species-level taxonomic concept label is sufficient to indicate the record’s identity with provenance from an expert-authored taxonomy. A full representation of that source’s taxonomic signal, and of its conflicts with other published signals, should live outside of the occurrence record.

In summary, the basic design preference for DwC over TCS and the way in which the former is typically implemented, are choices that compromise the taxonomic coherence of biodiversity data packages being processed for aggregation.

## Aggregation and authorship

Some aggregators may disagree that systematic syntheses are novel theories, or challenge the significance that we ascribe to their pragmatic creation. Several recent (self-)assessments of aggregators do not focus on the issue of *backbones* as an important challenge they must own and overcome (19-21, 33, 83). Instead, we find the following position symptomatic (19: 73; italics added for emphasis): “*Agencies* such as the ALA and GBIF *enable* observations *to be recorded directly* to their systems. These records are reviewed before being ‘published’, but the ALA and GBIF are *not* the data provider and therefore *cannot assume responsibility* for these records”.

In contrast, we argue that by promoting backbones as the primary means of offering content, aggregators act not just as data access facilitators but as data identity authors. It is not equitable to suggest that these issues can be dealt with “at the source”, or that “data errors are best addressed through collaboration between all relevant agencies” (19: 73), when the entity at the top holds all the cards.

Furthermore, aggregators frequently publish their syntheses under conventions that obscure individual expert authorship. This is not an all-or-nothing phenomenon: often there are linkages to individual authors or authored-attributed sources at lower-level nodes of the synthesis. Yet for instance, the 50+ sources for the GBIF taxonomy are all cited as initiatives, i.e., institutionalized catalogues, checklists, databases, species files projects, etc. No individual authors are named ‘on the surface’, i.e., on the top-level homepage. The entire synthesis is published by the “GBIF Secretariat” (65). The Open Tree of Life project (62, 73, 84), which is groundbreaking in its ability to accommodate individual source publications, nevertheless publishes its periodical synthesis versions without naming individual authors. The World Register of Marine Species lists ca. 260 taxonomic editors, making an effort to accredit views to editors and primary publications at lower levels. That said, the “WoRMS Editorial Board” (67) publishes the entire synthesis.

Unlike just three decades ago, aggregators now have technical means to appropriate and synthesize thousands of expert-authored monographic, revisionary, and phylogenetic publications into novel syntheses that *on the surface* accredit only an initiative or committee. However, the social costs of creating syntheses with obscured author accreditation can be considerable. To expert authors, this may signal not only that some low-level form of intellectual appropriation is taking place (85), but more disruptively, that accountability standards for the process of generating new systematic theories are shifting. For any particular taxonomic group, the synthesis may well represent the majority view. But who is responsible for defending that view scientifically? Who can be engaged to receive credit, and also criticism, for creating the synthesis as a unique hypothesis of how the natural world is structured?

Historically, the field of systematics has provided built-in opportunities for individual authors to reaffirm, expand, or challenge existing classifications through persistent publication venues. If aggregators publish syntheses that are not just novel but authorless in the conventional sense, they are thereby redesigning both the content of systematic theories and the social mechanisms for assigning responsibility for such content. Particularly the latter action constitutes a change in the power structure between aggregators and individual experts, to the experts' detriment.

## Consequences of disenfranchising taxonomic experts

Taxonomic perspectives that conflict with an aggregator’s synthesis have no equitable way to compete in the same environment. Consider, for instance, the effect of this single-party system on the aggregators' relationships with early-career systematists. Many systematists will find their path into the field because of the following circumstances: they have a passion for understanding biodiversity and have likely had a formative experience that existing systematic hypotheses for a particular lineage are empirically inadequate. *Not* accepting the consensus is an important motivator for systematic careers. The mission to revise challenging groups solidifies the intellectual identity of systematists, who must become professional consensus disruptors to make their mark in science.

Systematic challenges that persist today are not trivial. Resolving them requires narrow specialization; hence the term expert is appropriate to identify an individual’s record of training and accomplishment. The research products of early-career systematists, including monographic or revisionary treatments generated to meet thesis requirements, are usually published in international, peer-reviewed journals. Traditionally, this is how novel, high-quality hierarchies become part of the systematic knowledge base while advancing systematists’ career trajectories.

It should be unproblematic for a graduate student’s published monograph, including the therein newly identified occurrence records, to *immediately* be integrated with global aggregator environments – Later that month, the records were;ven and especially if the new classification conflicts with the aggregator’s synthesis, as it almost invariably will. Technical and social barriers to representing these data *as published* have a strong antagonistic effect. Evidently the monograph was good enough to earn a doctoral degree and peer-reviewed publication. However, propagating the same signal within the aggregator environment requires another level of validation that can appear unattainable.

The example of the monograph of *Zelus* Fabricius, 1803 sec. Zhang et al. (86) illustrates how the current power structure can operate. This publication accommodates 71 species-level concepts, including 24 new species names and numerous other taxonomic and nomenclatural changes. The monograph also identifies 10,628 specimen records. These can be downloaded in DwC-compatible format from the article website of the *Biodiversity Data Journal* (87). The example is fair precisely because the occurrence records were shared openly at the time of publication and in accordance with conventions that the aggregator endorses (88). Hence, this is not a case of erroneous execution.

Zhang et al. (86) was published in early July, 2016. Later that month, the records were aggregated in GBIF as well. However, the representation conventions *newly* constrained the original set of occurrence records in at least two critical ways. First, the aggregator (88) showed only 409 of the source records (3.8%), i.e., those cited in the publication’s main text. This constraint was applied by the journal publisher. Second, for the duration of nearly 220 days – i.e., until the next GBIF Backbone update took place (89) – the aggregator validated only 17 of 78 taxonomic names (21.8%), leaving 61 species-level epithets unrecognized. Among other consequences, this meant that the epithets of 40 holotype specimens referenced in Zhang et al. (86) were changed to show only the genus-level name “*Zelus* Fabricius, 1802” in the GBIF aggregate (though see above: the epithets were retrievable for individual records and full downloads). The timing of this update was solely controlled by the aggregator.

The GBIF Backbone is not open for direct edits by expert authors. Furthermore, directly submitting (e.g.) a Darwin Core Archive data file (90) can be challenging for individual authors. Experts can only submit “issues” for tracking through the ChecklistBank code repository (72), with the expectation that someone with high-level access rights – though typically less specialized taxonomic expertise – will address them (22).

It is clearly positive that within seven months, the GBIF records’ identifications were adjusted to reflect Zhang et al. (86). At the same time, neither that event nor the entire process can be said to have empowered the authors of the monograph. The vibe that experts get from an aggregator is more like this: *we* choose if we want to represent your knowledge, and when. One might even say: if an aggregator wanted to harm early-career systematists where it hurts their academic identities the most, controlling the form and content of their thesis-derived occurrence record identifications and systematic hypotheses is unfortunately quite effective. In analogy to the effect of negative on-line reviews for contractors, the experience of feeling disenfranchised will endure long after the records are adjusted.

## Backbone-based data signal distortion: an example (Figure 1)

Organismal groups like plants and birds are revised so frequently that multiple incongruent systems remain in use *simultaneously* (30-31, 78, 82, 91). In some instances, the parallel existence of regional biodiversity data cultures that endorse conflicting taxonomies reflects an equilibrium condition (82). Persistent conflict is also common in phylogenomics, where competing tree hypotheses can appear in short order and continue to attract endorsements by third-party researchers (92).

**Figure 1.**
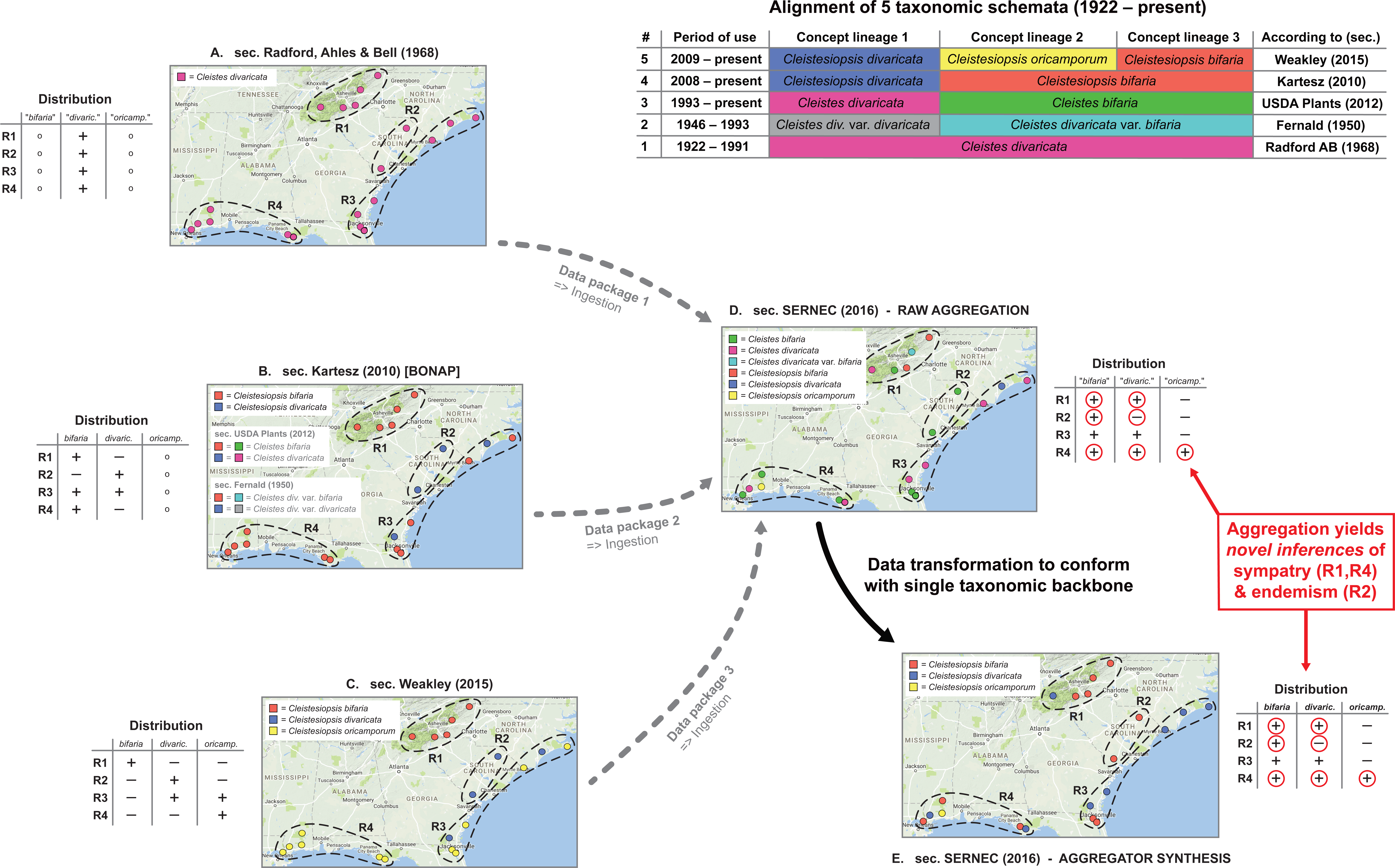
Backbone-based aggregation disrupts coherent biodiversity data packages. 'Most real' example adopted from Franz et al. (30). The top right table presents an alignment of five different taxonomies for the *Cleistes/Cleistesiopsis* complex sec. Radford et al. (101), Fernald (102), USDA Plants (103), Kartesz (104), and Weakley (94). Columns indicate the relative congruence between different taxonomic concepts, whereas rows show the period of usage, validly recognized names, and sources. A-E: five representations of the same set of 20 specimens provided by the SERNEC Data Portal (93), with distribution maps that identify four ecoregions R1-R4 (right), and tables displaying the ecoregion-specific presence (+), absence (–), or inapplicability (o - i.e., name not available) of occurrences identified to taxonomic concept labels. A-C: concept occurrence patterns according three reciprocally incongruent, yet internally coherent taxonomies; D: raw (unprocessed) aggregate of A-C, where each source contributes a complementary subset (data package) of the 20 specimens – hence six taxonomic names are shown; E: backbone-based transformation of D. Both D and E support new biological inferences (red circles) regarding the sympatry of multiple entities of the complex in ecoregions R1 and R4 (= false positives), and the local endemism of an entity labeled *bifaria* in R2 (= false negative), which is possible if pro parte synonymy relationships are not coherently transposed in the backbone-based synthesis.

Taxonomic backbones are deeply problematic in such situations. Our example of Figure 1 shows how a backbone may distort the identities of occurrence records *qua* aggregation, and thereby support inferences that conflict with every source-derived data signal. The example is a simplification of figures 1 and 2 in Franz et al. (30), showing only 20 occurrences that originate from the Southeast Regional Network of Expertise and Collections herbarium portal (93). These were deliberately filtered to illustrate the effect.

The starting conditions are common enough. A group of endangered orchids is subject to multiple taxonomic revisions – reviewed in Weakley (94) – over a relatively short time interval. The revisions have led to a complex matrix of name usage relationships between different sources, with incongruent perspectives regarding species-level concept granularity and genus-level concept assignment. In spite of this, all expert-promoted views (A-C in Figure 1) concur internally that two of the four ecoregions herein identified – i.e., R1 and R4 – do not harbor individuals from multiple recognized species in the group. Furthermore, no expert author has ever recognized the presence of an entity with the epithet *bifaria* in the ecoregion R2. That said, and depending on the taxonomies being aligned, the names *bifaria* and *divaricata* have varying, pro parte synonymy relationships.

Backbone-based aggregation fails to preserve the expert-sourced taxonomic coherence of the 20 occurrence records in our example (D and E in Figure 1). The data-transforming process yields novel biological inferences with no support from any input source; viz. R1 and R4 are regions of sympatry for the orchid group, whereas region R2 harbors an entity labeled *bifaria* yet not *divaricata*. In other words, the interaction of taxonomic pluralism at the source level with backbone-based aggregation can create type I or type II errors in distributional signals (i.e., false positives or negatives) for which only aggregators may claim responsibility. Given the power balance inherent in the aggregation design, such errors need not be frequent to have a chilling effect on biologists’ trust.

## An objection

Two reviewers put forward an important criticism that we believe deserves special discussion. The reviewers suggested that no self-respecting biodiversity researcher would or should trust aggregated data blindly, that indeed careful data cleaning is almost always necessary and expected to render the downloaded data fit for purpose, and that therefore aggregator services need to be understood mainly or merely as data discovery tools. We reject this deflationary view for four reasons. For one, aggregators frequently blur the lines between advertising their services just as a data discovery tool or as a more powerful data signal tool. Second, the biases inherent in using unitary backbones remain in place even if users are only interested in discovering all relevant data for their research purpose. If the backbone-based data record modulations are not easily retrievable through primary on-line interfaces, then users are significantly constrained in their ability to design search queries with high rates of precision and recall (81). Third, let us assume that labor-intensive off-line data quality review and correction efforts are indeed the norm, prior to publishing. Then why must the fruits of these efforts remain *outside* of the aggregator’s environment? Why can they not immediately flow back into the same aggregation domain, while recording the provenance of expert changes? In other words, if the workflow of rendering data fit for purpose flows only in one direction, i.e. from the on-line aggregate to the off-line quality review and publication, then our criticism of the design stands.

Lastly, while such external quality review and cleaning routines can be achieved by taxonomically trained biodiversity informatics experts, we may not expect that non-specialists will have the ability to undertake such actions, both intellectually and also technically. This is why they should be relocated and accredited inside the environment.

A helpful analogy is a home owner who gives a contractor a blueprint specification (probably incomplete) for a new kitchen. The owner then discovers that the contractor has built a different kitchen that could in principle be disassembled and rebuilt by the home owner to the original specification. The contractor points out that each piece of building material retains its original labeling as assigned in the blueprint. The home owner *might* be able to understand and reconstruct what the contractor did, if the owner can come by adequate training and resources. Nonetheless, this in-principle possibility misses the point. If the contractor is not going to build according to the desired blueprint for reasons of practicality, it would at least help to follow a modular design so that the home owner can easily reorganize on the fly with minimal effort and training. Similarly, the taxonomic provenance and enacted expert validation efforts must be an explicit part of aggregated biodiversity data packages.

## Conclusion

Our diagnosis implies a clear set of recommendations for moving forward. A first step is to recognize that trust is not just a feature of data signal quality but also a consequence of the social design of aggregation and the resulting power balance between data contributors and aggregators. This insight should not count as a justification for contributors to withhold occurrence records (26, 33). However, declining to trust an out-of-balance social design *does* make sense in light of well-established thinking about cooperative knowledge systems (48-51, 55, 57). Aggregators are well advised to accept an inclusive notion of trust that gives weight to equitable social engineering.

A second step is to acknowledge the accountability gap created by downplaying the impact of taxonomic backbones. Providing a clearinghouse for occurrence records is not simply a matter of cleaning up records provided by source collections. Instead, one needs a way to collate those records into meaningful biological datasets for swathes of life where no universal, consensus hierarchy exists. The practical need for such a hierarchy does not erase its scientific status as a classification theory (42).

We suggest that aggregators must either author these classification theories in the same ways that experts author systematic monographs, or stop generating and imposing them onto incoming data source representations. The former strategy is likely more viable in the short term, but the latter is the best long-term model for accrediting individual expert contributions. Instead of creating hierarchies they would rather not 'own' anyway, aggregators could provide services and incentives for ingesting, citing, and aligning expert-sourced taxonomies (30).

As a social model, the notion of backbones (2) was misguided from the beginning. The idea disenfranchises systematists who are by necessity consensus-breakers, and distort the coherence of biodiversity data packages that reflect regionally endorsed taxonomic views. Henceforth, backbone-based designs should be regarded as an impediment to trustworthy aggregation, to be replaced as quickly and comprehensively as possible. We realize that just saying this will not make backbones disappear. However, accepting this conclusion counts as a step towards regaining accountability.

A third step is to refrain from defending backbones as the only pragmatic option for aggregators (95). The default argument, also expressed by some reviewers, points to the vast scale of global aggregation while suggesting that only backbones can operate at that scale now. The argument appears valid on the surface, i.e., the scale *is* immense and resources are limited. Yet using scale as an obstacle is only effective *if* experts were immediately (and unreasonably) demanding a fully functional, all data-encompassing alternative. If on the other hand experts are looking for *token actions* towards changing the social model, then an aggregator’s pursuit of smaller-scale solutions is more important than succeeding with the ‘moonshot’. And clearly, such solutions *have* been developed, often under the term “taxonomic concept approach” (79, 80, 82, 96-98). Avibase (82) uses this approach to manage more than 1.5 million taxonomic concept labels from 150+ checklists published over 125 years. This scale, covering some 10,000 avian species-level concepts as currently recognized by distinct sources, was achieved largely through an exceptional single-person effort. Aggregators seeking to improve expert trust are therefore better advised to embrace such incremental design improvements now rather than focus too much on global scalability as part of a flawed all-or-nothing argument. Otherwise, what is sold as pragmatism begins to sound dogmatic. In the presence of successful medium-sized solutions, the unwillingness to adjust designs – even for pilot projects – will likely be viewed by the expert community as a strategy to double down on the current power structure.

Another seemingly pragmatic argument is that many data packages to be aggregated are themselves syntactically underspecified. Again, while true on the surface, this does not preclude the design of pilot systems that enforce TCS-over DwC-based syntax – at a scale commensurate with individual expert publications (99). Failures to propagate such innovations at greater scales, and continued preferences for Darwin Core for purposes beyond its design scope, send the wrong message to experts.

Which brings us to the final conclusions. We intended to diagnose a systemic impediment to trusting biodiversity data aggregation. Whenever we mentioned specific aggregators, our purpose was solely to exemplify a broader design paradigm. While we cannot claim to have a full-scale solution on hand, the key directive is to develop new technical pathways and social incentives for experts to contribute directly to the validation of taxonomically coherent data packages in a greater aggregate. This will require new, broad-based political will in order to ensure its priority of the agendas of aggregators.

Over the past several decades, biodiversity data aggregation has taken a turn away from the agenda of promoting the careers of individual systematists. While recognizing the reasons aggregators had for taking this path, we suggest that the price of doing so, translated into declining trust, is higher than aggregators may have expected. The price is likely also higher than aggregators can afford in order to maintain long-term viability in the biodiversity data ‘economy’.

We view this diagnosis as a call to action for both the systematics and the aggregator communities to reengage with each other. For instance, the leadership constellation and informatics research agenda of entities such as GBIF or Biodiversity Information Standards (100) should strongly coincide with the mission to promote early-stage systematist careers. That this is not the case now is unfortunate for aggregators, who are thereby losing credibility. It is also a failure of the systematics community to advocate effectively for its role in the biodiversity informatics domain. Our aggregation designs mirror how much we value individual expertise. Shifting the power balance back to experts is therefore a shared interest.

## Acknowledgments

The authors thank the four referees for their constructive and detailed feedback. We are also grateful to Erin Barringer-Sterner, Andrew Johnston, Jonathan Rees, David Remsen, David Shorthouse, and Guanyang Zhang for helpful discussions on this subject.

